# Which pLM to choose?

**DOI:** 10.1101/2025.10.30.685515

**Authors:** Tobias Senoner, Ivan Koludarov, Joshua Günther, Amarda Shehu, Burkhard Rost, Yana Bromberg

## Abstract

Protein-language models (pLMs) provide a novel means for mapping the protein space. Which of these new maps best advances specific biological analyses, however, is not obvious. To elucidate the principles of model selection, we benchmarked fourteen pLMs, spanning several orders of magnitude in parameter count, across a hundred million protein pairs, to assess how well they capture sequence, structure, and function similarity. For each model, we distinguish *inherent information*, i.e. signal recoverable from raw-embedding distances, and *extractable information*, i.e. signal revealed through additional supervised training.

Three key results emerge. First, pLM protein representation space is inherently different from the space of biological protein representations, i.e. sequences or structures. Here, a size-performance paradox is salient – *mid-scale* foundation models are as good as much larger ones in reflecting all tested biological properties. Second, pLM representations compress and store biological information in proportion to model size. That is, a lightweight feed-forward network can be trained on embedding pairs to predict said biological properties well – a capacity dividend. Finally, we observe that a task-specific learning radically reshapes the embedding space, gaining inherent understanding of the task, but garbling any further extractions.

In other words, smaller pLMs can provide efficient and compute-light general insight. Larger models are advantageous only when fine-tuning is planned to accomplish a specific task. Furthermore, representations generated by “specialist” models are not immediately generalizable throughout protein biology. Thus, for pLMs, bigger isn’t always better.

## Introduction

Different protein language models (<u>pLMs</u> – term introduced in ProtTrans (Elnaggar et al., 2022)) can be conceptualized as distinct maps of the protein sequence space. Just as different cartographic projections emphasize different geographical features, different pLMs construct fundamentally different representations of the same biological landscape through their learned protein embeddings. With the increasing diversity of pLMs, (Heinzinger & Rost, 2025; Wang et al., 2025; Weissenow et al., 2022) a critical challenge has emerged: which pLM to choose for a specific biological application?

To answer this question, we distinguish between two types of information encoded within pLM embeddings: (1) **inherent information**, which is directly accessible through simple arithmetic computation on raw embedding vectors, e.g. to establish protein distances/similarities and (2) **extractable information**, which requires subsequent embedding-based supervised machine learning, e.g. to predict specific protein features. Understanding this distinction reveals different model capabilities and computational trade-offs.

Current pLMs span diverse architectures, scales, and targets: from ESM2 (Lin, Akin, Rao, Hie, et al., 2023) and ProtT5 (Elnaggar et al., 2022) models of eight million (8M) to 15 billion (15B) free parameters to specialized models, such as CLEAN, targeting enzyme labeling,(Yu et al., 2023) and ProtTucker, targeting structural domain classification (Heinzinger et al., 2022). Unlike in natural language processing (NLP), where larger models typically yield better results (Hoffmann et al., 2022), the relationship between the model size, i.e. number of free/trainable parameters, and protein representation quality remains unclear (Schmirler et al., 2024; Teufel et al., 2022).

How does one pick a pLM, a map, that is most representative of the (protein) world? For geographical maps, this selection is not usually done in isolation, but rather with focus on their usefulness for a specific task. Note that task-relevant features can we added to any map, but this would, trivially, change the map, even as it becomes more suitable for that specific task. Evaluating this new map is then different from judging the quality of the original. Evaluating pLMs similarly requires use-cases, i.e. a biologically meaningful task that can be completed using embeddings directly and without further training, i.e. changing the representation/map.

Here, we set out to evaluate pLM quality by relying on a long-standing computational biology problem of identifying protein function – a task plagued by multidimensionality and conflicting annotations (Ashburner et al., 2000; Gillis & Pavlidis, 2013; Radivojac et al., 2013; Rembeza & Engqvist, 2021; Schnoes et al., 2009). In the absence of complete and precise functional labels, we fell back on the practice of “function transfer by homology”, where protein pairs deemed related, can be assigned shared function. That is, we evaluated protein functional similarity using the principle that it’s easier to state “these proteins have the same function” (Mahlich et al., 2018; Prabakaran & Bromberg, 2025a) than to define function individually. Incidentally, this approach allows for direct evaluations of protein projections in the pLM space, i.e. functionally similar proteins should ideally be close on this map.

To this end we systematically compared representations generated by 14 commonly used pLMs. In addition to protein functional similarity, measured by their HFSP score (Mahlich et al., 2018), we assessed two controls: sequence similarity, reported as the percentage pairwise sequence identity (PIDE), and structural similarity, reported by their TM-score (Y. Zhang & Skolnick, 2005). Specifically, we asked whether embeddings could capture sequence similarity well and, more importantly, whether this feature is the *only* one that embeddings would capture; the latter outcome would be the worst-case scenario for pLM use, where model representations would not contribute novelty to our understanding of proteins. Structure similarity was our second, more advanced, baseline, where pLM success would suggest that model representations are able to capture molecule biophysical features – a likely outcome, given earlier success of sequence-based structure prediction models. As sequence, structure, and function similarity are correlated but not identical, we expected to observed differences in model performance for each of these values.

By analyzing both inherent information (raw embedding distances) and extractable information (supervised learning performance), we addressed two key questions: **“Does pLM embedding similarity inherently capture biologically meaningful protein relationships?”** and **“Do larger pLMs necessarily create better protein representations?”**

Our findings reveal that all pLMs retain more extractable information than is inherently available. Moreover, inherent information is similar across larger and smaller models of the same family, with benefit plateauing early relative to model size. Access to extractable information, however, is significantly improved in larger models. Importantly, task-specific training dramatically reshapes the embedding spaces, changing the map to create specialized and significantly less generalizable representations.

## Methods

### Data preparation

To evaluate the performance and generalizability of protein language models (pLMs), we prepared two datasets. Our primary dataset, **SwissProt-pre2024**, was constructed for model training, validation, and testing from the Swiss-Prot database release 2024_01 (The UniProt Consortium et al., 2025). To create non-redundant data splits, we first partitioned all proteins using the *cluster* mode of MMseqs2 (Steinegger & Söding, 2017). Clusters were defined by a minimum of 30% pairwise sequence identity (--min-seq-id 0.3) and 80% coverage of the shorter sequence (-c 0.8,--cov-mode 1), using a sensitivity setting of -s 7.5. The resulting clusters were then split in a 70/15/15 ratio for training, validation, and testing, respectively, stratified by cluster size to ensure a balanced distribution. This procedure yielded a training set of 375,209, a validation set of 86,546, and a test set of 80,514 sequence non-redundant proteins.

To assess model generalizability on novel proteins, we constructed a second, more stringent test set named **New2024**. This set comprises proteins that are not only recently reported but also sequence-dissimilar to our primary dataset, based on UniRef50 cluster membership. We identified all UniRef50 clusters present in the 2025_01 release but absent in the 2024_01 release. From this set of new clusters, we established a “truly novel” subset by applying two strict filters. First, we retained only those clusters where none of the members appeared in *any* UniRef50 cluster from the 2024_01 release, ensuring that they represent new sequences rather than re-clustered known ones. Second, we filtered out any remaining clusters containing UniParc entries to exclude any sequences that were previously available in public databases. We then compiled all sequences from the remaining clusters and filtered for those with high-confidence Protein Existence (PE) annotations (level=1, i.e. “evidence at protein level” or level=2, “evidence at transcript level”) to ensure that we retained only experimentally validated sequences. The final **New2024** set was formed by intersecting this high-confidence candidate pool with the members of our “truly novel” clusters. This rigorous filtering process yielded a final test set of 1,237 proteins from 833 distinct, truly novel UniRef50 clusters.

### Protein similarity metrics

To establish the ground truth for model evaluations, we computed three distinct similarity metrics for protein pairs within all our datasets: sequence identity, structural similarity, and functional similarity. **Sequence Similarity:** we performed an all-against-all search using MMseqs2 (Steinegger & Söding, 2017). The search was configured as a high-sensitivity profile search with three iterations *(-s 7*.*5, --num-iterations 3, --e-profile 1e-10*). We generated full alignments (*-a, --alignment-mode 3*) for up to 1,000 target sequences *(--max-seqs 1000*) meeting an E-value cutoff of 0.001 *(-e 0*.*001*). From the resulting alignments, we extracted the fractional identity (fident), defined as the number of identical aligned residues divided by the total length of the alignment. We name this metric *PIDE* (percent pairwise sequence identity) for the rest of this manuscript. **Structural Similarity** was quantified using the *alntmscore* computed using Foldseek (van Kempen et al., 2024). We employed the identical search parameters as for sequence similarity computation. For the **SwissProt-pre2024** dataset, we used the pre-existing Foldcomp (Kim et al., 2023) compressed structures from the AlphaFold DB (v4) (Varadi et al., 2024). For proteins in the **New2024** set, which lack corresponding AlphaFold entries, we first predicted their structures using ColabFold (Mirdita et al., 2022). The structural alignment metric for all pairs is termed *TM-scor*e for the rest of the manuscript. **Functional Similarity** was assessed by computing the *HFSP* (homology-derived functional similarity of proteins) scores (Mahlich et al., 2018) from the MMseqs2 alignments above.

To ensure the quality of our ground-truth data, we filtered the resulting pairs based on established reliability thresholds for each metric (Figure S1). (1) For sequence similarity, we retained pairs only if they had a PIDE>0.3 and an alignment coverage of ≥80% for both proteins. (2) For structural similarity, we excluded any protein structure with an average prediction reliability (pLDDT) score≤70. We also required an alignment TM-score≥0.4 and an alignment coverage≥80% for both structures. (3) Finally, pairs were only considered functionally similar if they had a positive HFSP score (>0).

After filtering, the training set for further supervised learning, contained 379,566 proteins, resulting in: (1) 47,777,153 pairs of proteins with similar sequences, (2) 111,875,191 pairs with similar structure, and (3) 47,777,153 pairs with similar function. The validation set contained 8,447,555, 15,699,682, and 8,447,555 pairs, while the test set contained 8,731,253, 15,481,580, and 8,731,253 pairs, respectively. To equalize comparisons, we used this test set for all inherent or extractable evaluations in this work.

Note that this filtering step excluded the vast majority of protein pairs – the highly dissimilar ones, i.e. those with pairwise similarity too low to be detected by our methods. Thus, the analyses presented here primarily assess the capacity of embeddings to quantitatively capture measurable similarity, rather than to perform a binary discrimination between similar and dissimilar proteins (Figure S1).

### Protein language models (pLMs)

We analyzed 14 pLMs of different architectures, training approaches, and sizes. Foundation models included <u>ProtT5</u> (originally named ProtT5-XL-UniRef50; with 1.5B parameters; 1024-dimensional embeddings) (Elnaggar et al., 2022), <u>ESM-1b</u> (650M parameters; 1280-dimensional embeddings) (Rives et al., 2021), <u>ESM-2</u> models with 8M, 35M, 150M, 650M, and 3B parameters and 320, 480, 640, 1280, and 2560-dimensional embeddings respectively (Lin, Akin, Rao, Hie, et al., 2023), <u>Ankh</u> with 450M and 1.15B parameters and 768 and 1536-dimensional embeddings (Elnaggar et al., 2023), <u>ESM-C</u> models with 300M and 600M parameters and 960 and 1152-dimensional embeddings (Team ESM, 2024), and <u>ESM-3</u> with 1.4B parameters and 1536-dimensional embeddings (Hayes et al., 2025). Task-specific models included <u>CLEAN</u> (650M parameters; 128-dimensional embeddings), trained via contrastive learning for enzyme classification (Yu et al., 2023) and <u>ProtTucker</u> (1.5M parameters; 128-dimensional embeddings), optimized for CATH domain prediction through contrastive learning (Heinzinger et al., 2022).

We define a model “family” as a set of model variants developed for a single publication, sharing identical architectures and training procedures and differing only in parameter count and embedding dimensionality. For each protein sequence, we generated fixed-length embeddings by averaging per-residue representations from each pLM’s final hidden layer, producing protein-level embeddings suitable for pairwise comparison analyses.

### Model training and evaluation framework

We developed a comprehensive framework to quantify biological information encoded in pLM embeddings. Evaluations were conducted in two settings: (1) native embedding dimensions and (2) standardized 128-dimensional representations obtained via Principal Component Analysis (PCA) (Pearson, 1901).

For each setting, we employed two assessment approaches. First, as a non-trainable baseline, we computed Euclidean distances between embedding pairs. Second, we trained feed-forward networks (FFNs) with symmetric architecture: each n-dimensional embedding was processed through a hidden layer (n → 64), concatenated, and passed through successive layers (128 → 64 → 32 → 1) to generate predictions (Figure S2).

Models were trained independently to predict three biological properties: sequence identity (PIDE), structural similarity (TM-score), and functional similarity (HFSP score). In total, this yielded 180 models: 3 properties × 15 embedding sources (14 pLMs plus random control) × 4 approaches (native/PCA × Euclidean/FFN). Random control embeddings (1024 dimensions; generated from a standard normal distribution) established performance baselines. All models were implemented in PyTorch Lightning with consistent hyperparameters: learning rate 0.001, batch size 1024, maximum 100 epochs, and early stopping patience of 5 epochs.

Performance was evaluated using Pearson R^2^ correlation. Bootstrap analysis (1,000 samples) yielded 95% confidence intervals with standard errors <0.001 and are therefore not visible in the graphics.

To enable cross-model comparisons, we applied max-min normalization to all pairwise embedding distances, scaling each pLM to the range [0,1] via the transformation x_norm_ = (x – x_min_)/(x_max_ – x_min_), where x_min_ and x_max_ represent the minimum and maximum observed distances for that pLM. This normalization was applied prior to visual comparison in Figure 2 and for quantitative assessment via Wasserstein distance. The Wasserstein distance (W) quantifies the minimum “cost” of transforming one probability distribution into another (Villani, 2008). We computed it between normalized pairwise distance distributions to compare embedding space distributions across different pLMs.

## Results

We analyzed embedding spaces of 14 protein language models (pLMs, Table S1) across three protein similarity scores (PIDE, TM-score, and HFSP) and two embedding similarity modalities (Euclidean distance and supervised training). We primarily focused on the capacity of embeddings to quantitatively capture similarity, rather than to perform a binary discrimination between similar and dissimilar proteins. We discovered fundamental differences in inherent and extractable information present in these embeddings.

### Size-performance paradox for inherent information

We observed that representations of proteins, generated by pLMs, were inherently different from protein sequences or structures, i.e. molecule representations that currently serve as primary protein descriptors. Here, ***inherent information*** refers to the biological signal present in the *raw* protein embedding space, i.e. the distances between protein embedding pairs. To evaluate representations, we compared similarity of protein sequences to the similarity of their embeddings. Across all non-specialized pLMs, at best, the Pearson correlation R^2^ between protein sequence identity (PIDE) and Euclidean distance of embeddings was 0.51 (for ProtT5 embeddings; Table S1). Structural similarity (TM-score) was also poorly represented by embedding distances (R^2^=0.4 for Ankh-base). These observations strongly suggest that pLM-generated protein representations describe proteins in ways unlike sequence or structure alone.

Importantly, in evaluating the similarity of inherent information to protein-pair biological relationships across model families, we found that mid-size foundation pLMs matched or even outperformed larger models (Figure 1 and Table S1). For instance, within the ESM-2 family (Lin, Akin, Rao, & Hie, 2023), the 8M-parameter variant reflected functional similarity (Pearson correlation R^2^=0.28 with HFSP score) better than the 3B-parameter model (R^2^=0.26). The effect was qualitatively similar for sequence identity (e.g. ESM-2 8M R^2^=0.44 vs. ESM-2 3B R^2^=0.45) and structural similarity (e.g. ESM-2 35M R^2^=0.18 vs. ESM-2 3B R^2^=0.19). In the ESM-C family (Team ESM, 2024), the 300M parameter model outperformed its 600M parameter *sibling*, achieving higher Pearson R^2^ for sequence (PIDE= 0.49 vs. 0.47) and structure similarity (TM-score= 0.10 vs. 0.09), while matching performance for functional similarity (HFSP= 0.30 for both). We termed this counter-intuitive outcome the ***size-performance paradox*:** where adding parameters increases theoretical capacity but does not necessarily improve biologically meaningful organization of the inherent information, i.e. the *raw* embedding space.

**Figure 1.**
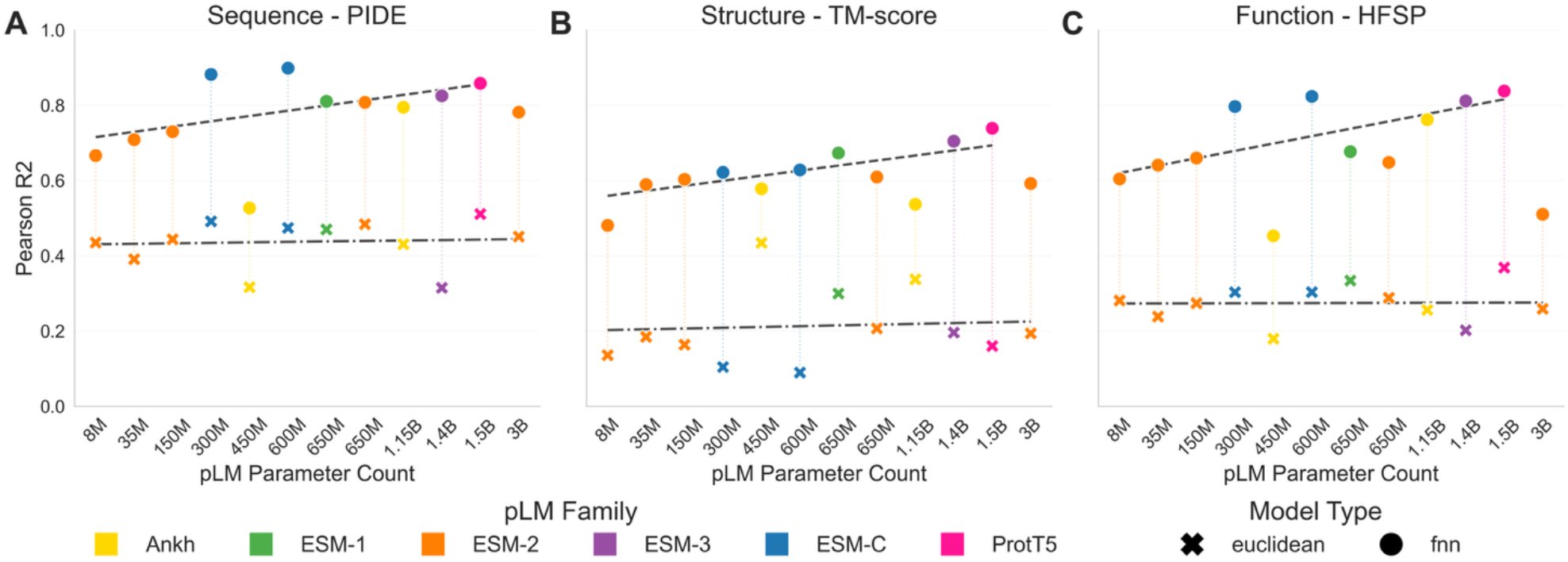
: Mid-scale non-specialist foundation pLMs produce the strongest “out-of-the-box” signal, while larger models are better with supervised tuning. Evaluated on SwissProt-pre2024 test pairs, for all models (x-axis, twelve non-specialist pLMs in order of their parameter count, colored by family), inherent embedding information (Euclidean distance of the two protein embeddings, marked by ×) is outperformed by additional supervised learning (prediction score from a feed-forward neural network, *FNN*, for the same embeddings, marked by •). Each panel reports the correlation (Pearson *R*^*2*^; y-axis) between the inherent information and the extracted information for one biological property of the pair: (A) sequence identity (PIDE), (B) structural similarity (TM-score), and (C) functional similarity (HFSP score). Trendlines across pLM parameter sizes (dashed line for FNN, dash-dot line for Euclidean) show that inherent information (×) remains flat with model size, with slopes of 0.0045, 0.0075, and 0.00087 for panels A, B, and C respectively, while extractable information (•) increases, with slopes 0.094, 0.09, and 0.13 (all values in change per billion parameters). Note: ESM2-3B was excluded from the FNN trendline fit as an outlier. Points (inherent and extractable information) of the same model are joined by dotted lines. All standard errors are below 0.001.

**Figure 2.**
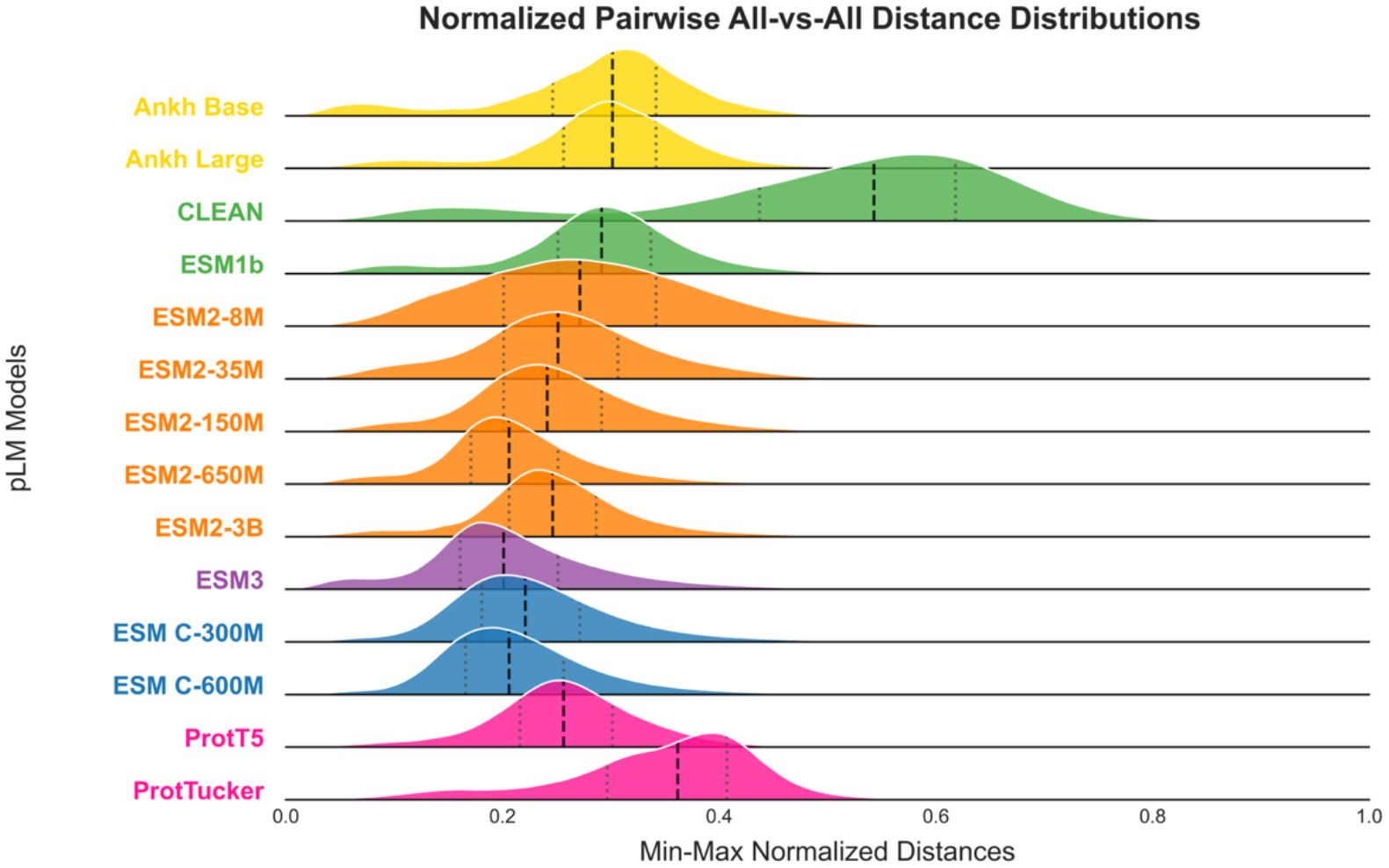
: Task-specific training distorts the protein-embedding space. Kernel-density ridges show the distribution of all*-*vs*-*all Euclidean distances between protein embeddings, after min–max normalization [0, 1] within each model. Rows are ordered by model family and within family by model size; colors match model names. Vertical lines indicate quartiles: dashed lines show medians, dotted lines show Q25 and Q75. Colors represent model families (shared by all family members): Ankh (yellow), ESM-1 (green), ESM-2 (orange), ESM-3 (purple), ESM-C (blue), and ProtT5 (pink). Foundation models ESM*-*2, Ankh, ProtT5, and ESM*-*C families display similar distance profiles within each family (e.g. ESM*-*2 variants share a common peak at ∼0.23). The task-specific CLEAN (green) and ProtTucker (Pink) are shifted to the right and are markedly broader than their parent ESM-1b or ProtT5, respectively.

Curiously, within the Ankh family (Elnaggar et al., 2023), the 450M parameter base model brought structurally similar pairs closer together than the 1.15B parameter large model (R^2^=0.43 vs. 0.34), but the smaller model was worse than the larger one for sequence and function comparisons. Indeed, in out-of-the-box comparisons across families, Ankh (base) outperformed all others in reflecting structure similarity, while ProtT5 was best among non-specialists at mapping sequence and function.

### Supervised learning reveals more information capacity in larger models

Inference of protein properties from sequence alone is fairly limited – while some features can be inferred from amino acid composition, models for predicting structure or function most often require sequence alignments to describe the evolutionary trajectories and sequence constraints (Jones et al., 2014; Rost & Sander, 1993). Protein embeddings, i.e. the end result of pLM training to capture sequence preferences in extant molecule collections, may inherently achieve similar ends. We thus further asked if pLM representations easily lend themselves to inference of protein characteristics. We assessed this ***extractable information*** captured by pLM embeddings by using these embeddings as input for supervised learning (shallow networks trained to predict one of: sequence identity, structure similarity, or shared function). Here we found that larger pLMs outperformed smaller ones, suggesting that the former capture a richer protein representation despite often weak inherent organization.

For example, networks trained with ESM C-600M embeddings surpassed ESM C-300M for extraction of all biological characteristics (PIDE Pearson R^2^=0.90 vs. R^2^=0.88, TM-score R^2^=0.63 vs. R^2^=0.62, and HFSP R^2^=0.82 vs. R^2^=0.80). Similarly, ESM-2 650M bested its smaller counterpart ESM-2 8M (HFSP R^2^=0.65 vs. R^2^=0.60, TM-score R^2^=0.61 vs. R^2^=0.48, and HFSP R^2^=0.81 vs. R^2^=0.67). However, the largest model in our set, ESM-2 with 3B parameters performed poorly, compared to its family’s smaller versions (PIDE R^2^=0.78, HFSP R^2^=0.51, and TM-score R^2^=0.59), suggesting a limit to performance improvement, perhaps given the current protein data availability. Again, Ankh behavior was somewhat of an outlier. Models trained on the 1.15B parameter embeddings were, as expected, better than the ones using the 450M parameter version for PIDE (R^2^= 0.79 vs. 0.53) and HFSP (R^2^= 0.76 vs. 0.45), but not for the TM-score (R^2^= 0.54 vs 0.58). Ankh behavior highlights the variability of large pLMs with even identical data sets and similar architectures.

Overall, performance generally improved with foundation model parameter count, regardless of architecture. Furthermore, all FNN models (extractable information) were better than Euclidean distance (inherent information) in reflecting all evaluated biological properties (highest Euclidian R^2^ vs. FNN R^2^; PIDE 0.53 ≤ R^2^ ≤ 0.9, TM-score 0.48 ≤ R^2^ ≤ 0.74, and HFSP 0.45 ≤ R^2^ ≤ 0.84). This observation highlights the necessity of further embedding processing when using protein representations instead of sequences. That is, embedding distances are not obviously constrained by evolutionary signals in the same way as protein sequences appear to be.

### Embedding dimensionality does not alter observed parameter-based trends

We evaluated embedding dimensionality across all models (ranging from 128 in CLEAN to 2,560 in ESM2-3B) as a factor in deciding performance. We standardized all embeddings to 128 dimensions using PCA (Methods) and repeated all experiments described above. Previous work in NLP showed that PCA-based dimensionality reduction can even improve the performance of pretrained embeddings while reducing computational costs (G. Zhang et al., 2024). Models trained on pLMs with more parameters still outperformed those trained on smaller variants (Figure S3 and Table S1).

### Task-specific fine-tuning reshapes embedding geometry

The three pLM families in this work exhibited consistent embedding space geometries (Figure S4). That is, pairwise embedding distance distributions between all proteins of a pLM tended to be more similar between pLMs of the same family than compared to other pLMs (Figure 2; Figure 3); higher Spearman correlations and lower Wasserstein distances within families vs. across families. For instance, ESM-2 variants (8M–3B parameters) were more similar than ESM-2 vs. Ankh, though smaller ESM-2 *siblings* had wider distributions (higher variance) than larger *siblings* (Figure 2).

**Figure 3.**
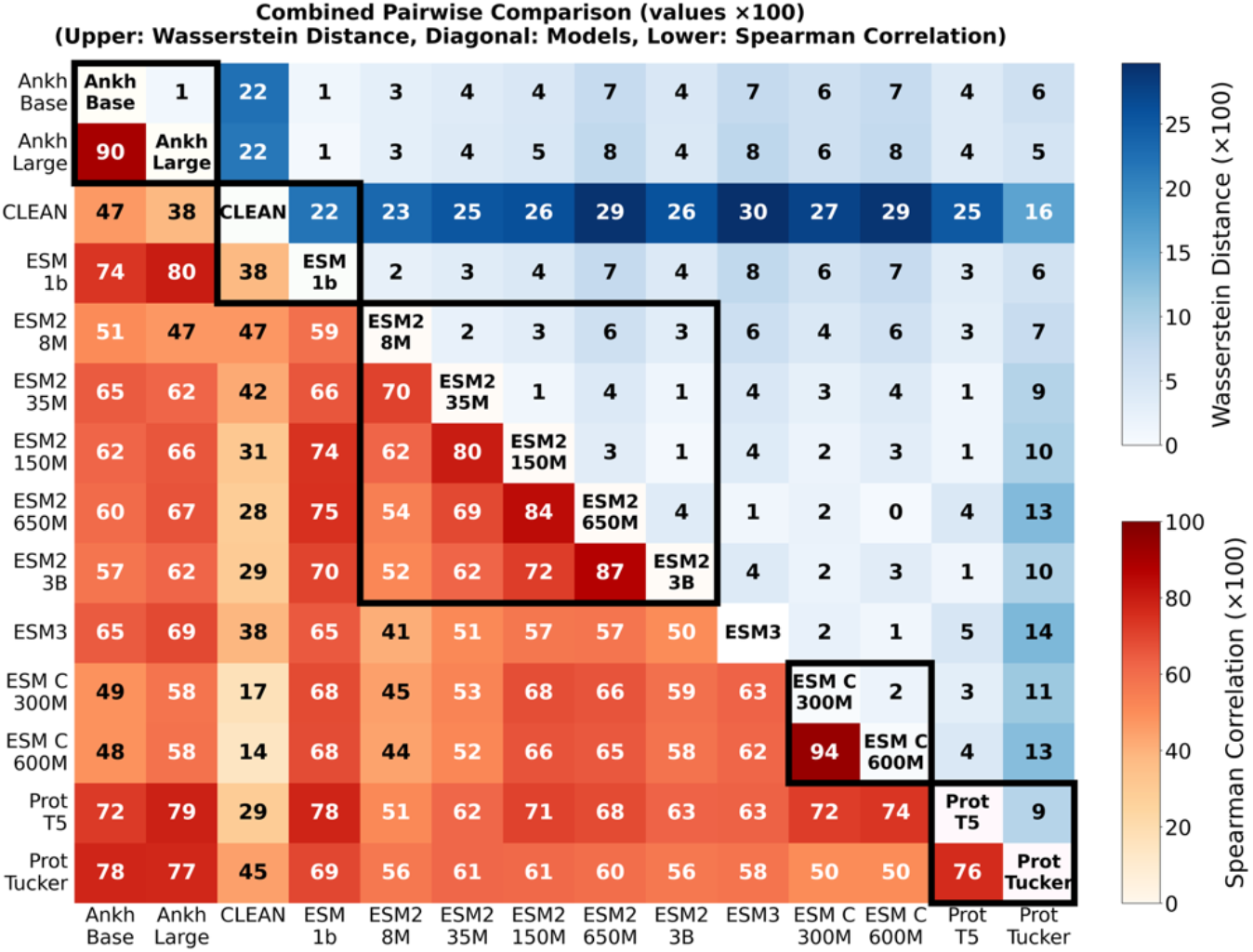
: Family identity and task-specific fine-tuning leave distinct fingerprints on protein-embedding geometry. For every protein pair in the SwissProt-pre2024 dataset, we computed the Euclidean distance between all protein embeddings produced by each of the fourteen pLM. The upper-right triangle (blue) presents normalized Wasserstein distances (W) between embedding-distance distributions for each pLM pair. For each pLM, its distribution of pairwise, raw-embedding distances was min–max normalized to [0, 1] and W computed to compare distribution shapes independent of scale. The lower-left triangle (orange) reports the Spearman rank correlations (ρ) between the model embedding-distance distributions, with cell coloring proportional to ρ. All values in the plot are displayed ×10^-2^ (e.g. 0.90 → 90). High ρ and low W values within model families – e.g. Ankh-base vs. Ankh-large (ρ=0.90; W=0.02) or ESM-C 300M vs. 600M (ρ=0.94; W=0.02) – highlight family-specific organization of embedding spaces. This similarity is lost for task-specific models; for example, CLEAN correlates poorly with its parent ESM1B (ρ=0.38; W=0.22).

In contrast, the embeddings inferred by supervised, contrastive learning models CLEAN and ProtTucker, displayed a substantially altered embedding space relative to their parents. CLEAN’s distribution was right-shifted (higher distances between protein pairs), bimodal, and broader than its origin ESM-1b, reflecting its objective of minimizing intra-class distances while maximizing inter-class separation for enzyme classification (Figure 2). This resulted in low similarity to ESM-1b ρ=0.37 and W=0.22 (Figure 3). ProtTucker – trained for CATH (Sillitoe et al., 2021) domain classification – distribution of embedding pairs also extended to the right, but maintained greater similarity to its parent ProtT5 (ρ=0.76, W=0.09). These findings demonstrate that foundation models within families implicitly preserve some aspects of their geometry, while task-specific objectives fundamentally reshape embedding geometry.

## Discussion

Through systematic analyses of protein language model (pLM) embedding spaces, we reveal fundamental trade-offs that challenge conventional assumptions about model scaling and highlight the effect of specialization on protein representations. These findings provide a framework for understanding when and why particular pLMs excel, guiding model selection for specific research needs. Although not considered in this work, we suggest that it may also be warranted to combine pLM embeddings on the basis of orthogonality (lack of correlation, Figure 3), thereby maximizing complementarity for further tuning.

### Size-performance paradox: diminishing returns for large pLMs

The counterintuitive observation that smaller foundation pLMs matched the performance of those (many) folds larger when exclusively using inherent information, departs sharply from the scaling behavior established in natural language processing (NLP). That is, we find that additional capacity yields no measurable gain in modelling protein relationships. For example, the ESM-C family, which has only 300M and 600M parameter variants, consistently outperformed earlier, larger models. In addition to the model’s architectural enhancements, this superior performance may stem from ESM-C’s expanded training data, which included UniRef (Suzek et al., 2015), MGnify (Richardson et al., 2023), and sequences from the Joint Genome Institute (Nordberg et al., 2014) clustered at 70% PIDE, making it substantially larger and, likely, more diverse than the datasets used by other models. If so, training data diversity, rather than model size, may be the key to advancing pLM performance.

The size-performance paradox reflects the fundamental difference between natural language and protein sequences. While natural language benefits from increasingly complex contextual relationships that larger models can capture, protein sequences may have more constrained and well-defined biological relationships that smaller models can encode directly. For current pLMs, and given the current relative paucity of data diversity (Koludarov, Senoner, unpublished data), the ∼300M parameter models may strike an optimal balance between representation usefulness and the inherent structure of protein sequence space.

We note that while our analysis has been limited by resource constraints barring the inclusion of even larger pLMs — a limitation shared in many academic environments — all results suggested that our identified patterns would hold true for them as well. Furthermore, among larger models in our set, superior performance of ProtT5 (1.5B) implied that alignment between model design, benchmarking objectives, and better-selected data, might be as (or more) decisive in ensuring overall model efficacy as parameter count.

### Capacity dividend: larger pLMs capture more extractable information

While smaller pLMs excelled in reflecting the inherent structure of the protein space, larger pLMs demonstrated superior capacity for extractable information, i.e. via subsequent supervised training on embeddings. This was particularly obvious for complex tasks like functional similarity prediction. For instance, the dramatic performance improvement of Ankh-1.15B over Ankh-450M with further training (R^2^=0.76 vs R^2^=0.45 for HFSP) indicated that larger pLMs encode more sophisticated representations, requiring additional effort to unlock.

This pattern suggests a fundamental trade-off in protein language modeling: immediate accessibility (inherent) versus ultimate potential (extractable). Larger models appear to encode biological information in more complex patterns that are less immediately interpretable but can be leveraged effectively through supervised training. This pattern has been observed even at the 100B parameter scale, where improved perplexity does not guarantee better downstream performance without task-specific fine-tuning (Chen et al., 2025). Yet the incremental nature of these improvements — even the largest models in this study struggled to exceed Pearson R^2^ of 0.74 and 0.84 for structural and functional similarity, respectively — suggests that current architectures may be approaching fundamental limitations in protein representation rather than simply requiring more parameters.

### Geometry-warp by task-specific training

The dramatic embedding space alterations observed for task-specific models, e.g. CLEAN (Yu et al., 2023), optimized to classify enzymes, and ProtTucker (Heinzinger et al., 2022), optimized to classify protein structure by CATH (Sillitoe et al., 2021), revealed both the power and limitations of specialized training.

Reduced correlation between CLEAN and its parent foundation pLM ESM-1b (ρ=0.38) illustrates how task-specific training fundamentally warps embedding spaces, optimizing for specific biological properties while losing sight of general protein relationships. CLEAN’s warping results in a broader distribution of pairwise distances (Figure 2) and limits the model’s ability to adapt to new tasks through supervised learning. Here, for example, CLEAN performed worse than ESM-1b (Pearson R^2^ PIDE 0.69 vs. 0.81, TM-score 0.52 vs. 0.67, HFSP 0.46 vs. 0.68) – a result due to performance measurements unconcerned with enzymes.

Both non-specialist models, ProtTucker and CLEAN, expand their embedding space relative to their parent models, ProtT5 and ESM-1b, respectively. We suspect that ProtTucker’s expansion improves CATH domain discrimination while potentially compressing other biological signals into narrower ranges. CLEAN’s expansion magnifies enzyme-class separations but dilutes signals for structural classification. Note that CLEAN’s ability to capture function is clear in its inherent relatively high correlation with HFSP. However, its embeddings do not lend themselves to further tuning (Table S1).

The geometry of an embedding space governs how every downstream method – nearest-neighbor search, clustering, or a thin supervised head – interprets relatedness. Because supervised training usually adds only a lightweight head on top of frozen embeddings, a strongly warped base geometry leaves little latent information to reuse, explaining each model’s weak cross-task transfer despite their hundreds of millions of parameters. That is, while task-specific training can achieve superior performance for targeted applications, it sacrifices versatility that makes foundation models valuable for diverse biological questions.

### Practical pLM selection and future directions

Our findings enable clear recommendations for pLM selection based on research requirements and computational constraints. For immediate biological insights requiring minimal computational overhead, i.e. using raw embeddings, mid-size foundation models are clearly sufficient. For highly specialized tasks with well-defined objectives, task-specific pLMs, regardless of their size, are needed; be aware that these models sacrifice general applicability for specialized performance. Note that, here, using LoRA fine-tuning to create smaller models specific for each problem (Schmirler et al., 2024), instead of large-scale re-training, is likely most efficient. Indeed, when planning additional fine-tuning for specific applications larger foundation models are warranted.

Overall, the modest performance improvements due to parameter scaling (Figure 1) suggests that current model limitations may stem from fundamental data and methodological constraints rather than from insufficient model capacity. That is, in addition to model size, the choice between pLM families should be driven by specific biological properties of interest, with different architectures exhibiting distinct strengths. Moreover, the underrepresentation of protein diversity in training datasets (Avasthi & York, 2024; Karsch-Mizrachi et al., 2018; Mora et al., 2011) likely constrains all models regardless of size. Systematic biases in protein representations identified in recent work (Ding & Steinhardt, 2024; Marquet et al., 2024) further compound these limitations, suggesting that addressing data quality and diversity may yield greater improvements than scaling model size.

Future progress in protein language modeling thus requires fundamental shifts in approach. First, developing methods to better preserve and extract biological information from existing embedding spaces remains a significant challenge. A recent study exploring 13 different compression methods found that average pooling of residue representations to represent proteins remains superior (Vieira et al., 2025), however additional work in this direction is warranted given the difference between embedding-inherent vs. extractable information patterns identified herein. Second, acknowledging that not all embeddings of one pLM are created equal will lead to developments of more descriptive models. That is, if evaluation methods such as RNS (Prabakaran & Bromberg, 2025b) can identify embeddings of poorly captured proteins, should we train additional models specific to these poorly captured subsets or, rather, optimize generalized model training? Third, explicitly designing training approaches that optimize embedding spaces for multiple biological properties simultaneously, possibly through multi-task or carefully designed contrastive learning, may result in improved representations for a better understanding of the protein space. Finally, it is critical for any future progress to address the issues of data quality and diversity. Focusing on underrepresented protein families and taxonomic groups through strategic dataset expansion and improved data curation may address systematic bias in protein representations, leading to more robust pLMs – no petabytes of parameters needed.

## Conclusion

Our systematic comparison of 14 protein Language Models (pLMs) uncovered a two-tier effect of scale. For zero-shot use, the sweet spot for pLM size appears today in mid-scale foundation models. Adding billions of extra parameters does not improve this inherent information but consumes substantially more energy. Additional extractable information capacity dividend becomes clear through supervised training with a simple prediction head. Specialization is a double-edged sword: contrastive fine-tuning for enzymes (CLEAN) or CATH domains (ProtTucker) boosts performance on the target task, but it also warps the global geometry of the protein space thereby reducing model versatility. Taken together, the results argue for a task-driven model-selection strategy: use smaller foundation pLMs when rapid, low-cost insight is required; reach for the billion-parameter variants only when you can afford to fine-tune and need maximal ceiling performance; and deploy task-specific models solely when the biological question precisely matches their training objective. In short, judicious choice of pLM can save both computation and carbon without sacrificing accuracy, that is bigger is not automatically better.

## Supplementary Material

**Table S1:**
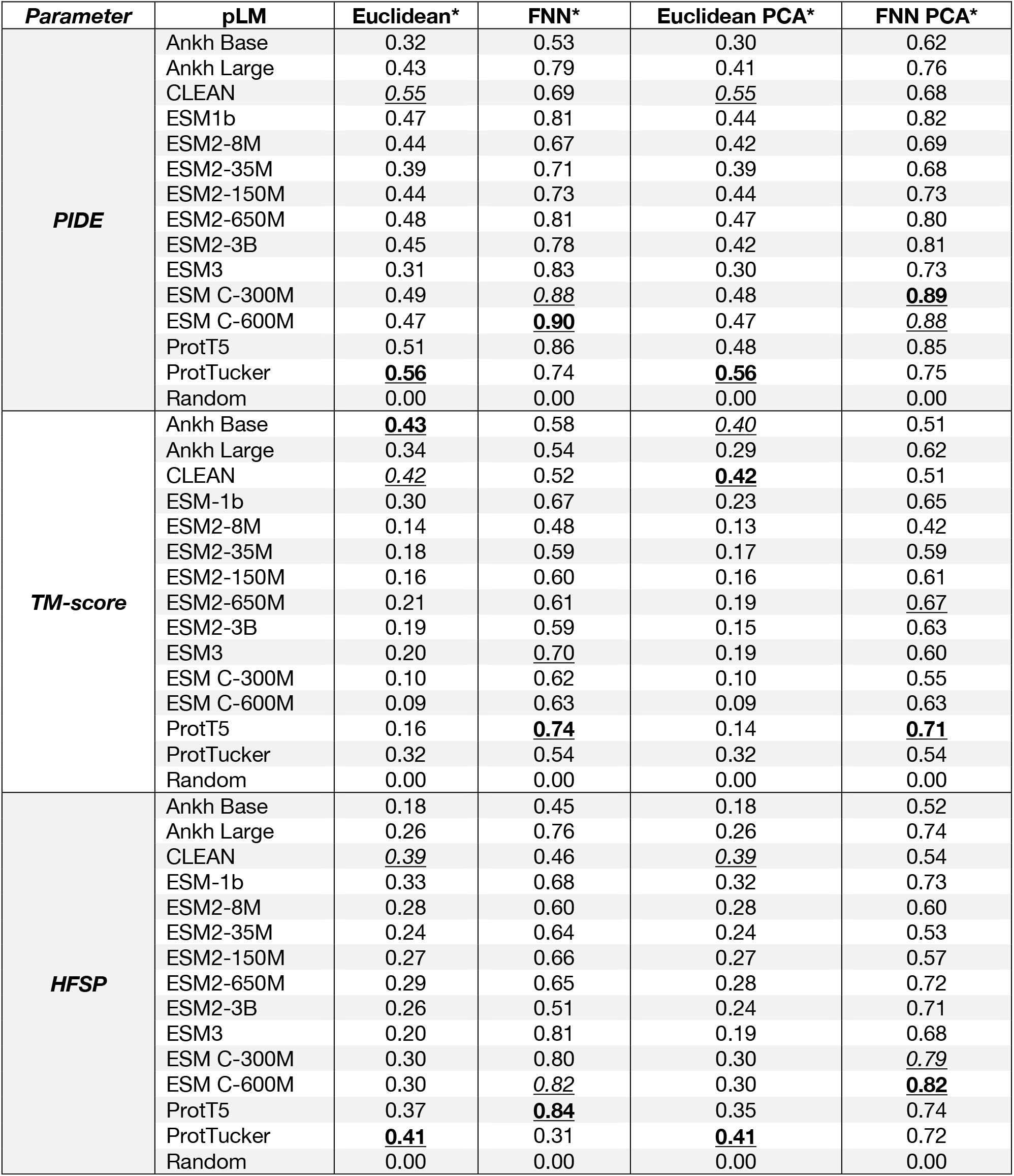
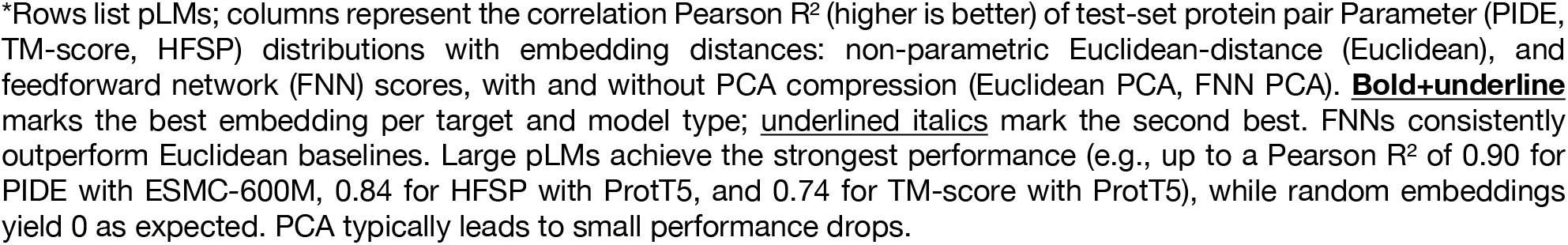
pLM performance in predicting pairwise PIDE, TM-score, and HFSP.

| *Parameter* | **pLM** | **Euclidean\*** | **FNN\*** | **Euclidean PCA\*** | **FNN PCA\*** |
| --- | --- | --- | --- | --- | --- |
| **PIDE** | Ankh Base | 0.32 | 0.53 | 0.30 | 0.62 |
|  | Ankh Large | 0.43 | 0.79 | 0.41 | 0.76 |
|  | CLEAN | <u>0.55</u> | 0.69 | <u>0.55</u> | 0.68 |
|  | ESM1b | 0.47 | 0.81 | 0.44 | 0.82 |
|  | ESM2-8M | 0.44 | 0.67 | 0.42 | 0.69 |
|  | ESM2-35M | 0.39 | 0.71 | 0.39 | 0.68 |
|  | ESM2-150M | 0.44 | 0.73 | 0.44 | 0.73 |
|  | ESM2-650M | 0.48 | 0.81 | 0.47 | 0.80 |
|  | ESM2-3B | 0.45 | 0.78 | 0.42 | 0.81 |
|  | ESM3 | 0.31 | 0.83 | 0.30 | 0.73 |
|  | ESM C-300M | 0.49 | <u>0.88</u> | 0.48 | **0.89** |
|  | ESM C-600M | 0.47 | **0.90** | 0.47 | <u>0.88</u> |
|  | ProtT5 | 0.51 | 0.86 | 0.48 | 0.85 |
|  | ProtTucker | **0.56** | 0.74 | **0.56** | 0.75 |
|  | Random | 0.00 | 0.00 | 0.00 | 0.00 |
| **TM-score** | Ankh Base | **0.43** | 0.58 | <u>0.40</u> | 0.51 |
|  | Ankh Large | 0.34 | 0.54 | 0.29 | 0.62 |
|  | CLEAN | <u>0.42</u> | 0.52 | **0.42** | 0.51 |
|  | ESM-1b | 0.30 | 0.67 | 0.23 | 0.65 |
|  | ESM2-8M | 0.14 | 0.48 | 0.13 | 0.42 |
|  | ESM2-35M | 0.18 | 0.59 | 0.17 | 0.59 |
|  | ESM2-150M | 0.16 | 0.60 | 0.16 | 0.61 |
|  | ESM2-650M | 0.21 | 0.61 | 0.19 | <u>0.67</u> |
|  | ESM2-3B | 0.19 | 0.59 | 0.15 | 0.63 |
|  | ESM3 | 0.20 | <u>0.70</u> | 0.19 | 0.60 |
|  | ESM C-300M | 0.10 | 0.62 | 0.10 | 0.55 |
|  | ESM C-600M | 0.09 | 0.63 | 0.09 | 0.63 |
|  | ProtT5 | 0.16 | **0.74** | 0.14 | **0.71** |
|  | ProtTucker | 0.32 | 0.54 | 0.32 | 0.54 |
|  | Random | 0.00 | 0.00 | 0.00 | 0.00 |
| **HFSP** | Ankh Base | 0.18 | 0.45 | 0.18 | 0.52 |
|  | Ankh Large | 0.26 | 0.76 | 0.26 | 0.74 |
|  | CLEAN | <u>0.39</u> | 0.46 | <u>0.39</u> | 0.54 |
|  | ESM-1b | 0.33 | 0.68 | 0.32 | 0.73 |
|  | ESM2-8M | 0.28 | 0.60 | 0.28 | 0.60 |
|  | ESM2-35M | 0.24 | 0.64 | 0.24 | 0.53 |
|  | ESM2-150M | 0.27 | 0.66 | 0.27 | 0.57 |
|  | ESM2-650M | 0.29 | 0.65 | 0.28 | 0.72 |
|  | ESM2-3B | 0.26 | 0.51 | 0.24 | 0.71 |
|  | ESM3 | 0.20 | 0.81 | 0.19 | 0.68 |
|  | ESM C-300M | 0.30 | 0.80 | 0.30 | <u>0.79</u> |
|  | ESM C-600M | 0.30 | <u>0.82</u> | 0.30 | **0.82** |
|  | ProtT5 | 0.37 | **0.84** | 0.35 | 0.74 |
|  | ProtTucker | **0.41** | 0.31 | **0.41** | 0.72 |
|  | Random | 0.00 | 0.00 | 0.00 | 0.00 |

\*Rows list pLMs; columns represent the correlation Pearson $R^2$ (higher is better) of test-set protein pair Parameter (PIDE, TM-score, HFSP) distributions with embedding distances: non-parametric Euclidean-distance (Euclidean), and feedforward network (FNN) scores, with and without PCA compression (Euclidean PCA, FNN PCA). **Bold+underline** marks the best embedding per target and model type; underlined italics mark the second best. FNNs consistently outperform Euclidean baselines. Large pLMs achieve the strongest performance (e.g., up to a Pearson $R^2$ of 0.90 for PIDE with ESMC-600M, 0.84 for HFSP with ProtT5, and 0.74 for TM-score with ProtT5), while random embeddings yield 0 as expected. PCA typically leads to small performance drops.

**Figure S1:**
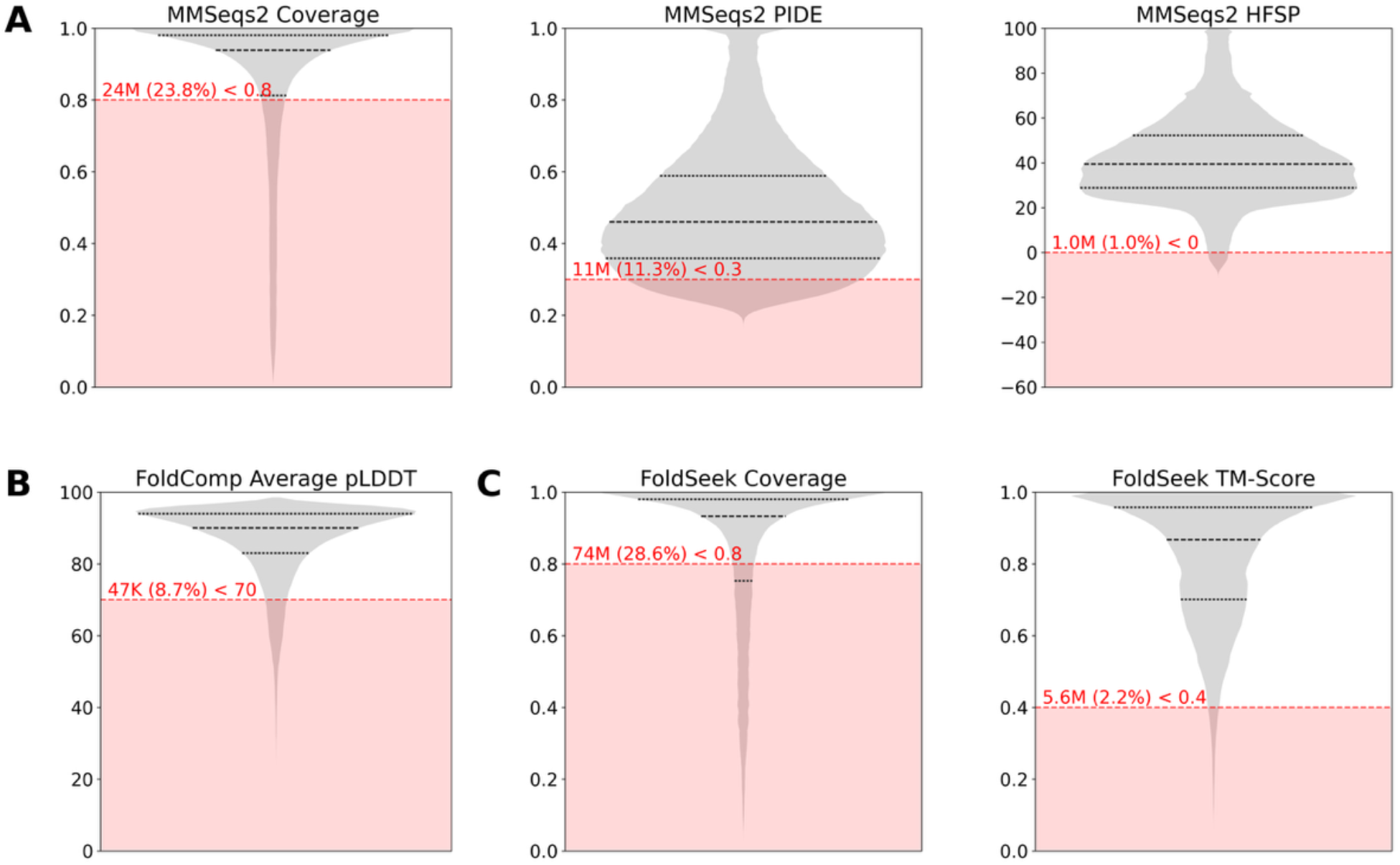
Distribution and filtering of protein similarity metrics. Violin plots illustrating the distribution of key quality and similarity metrics across protein pairs prior to filtering. Each plot visualizes the data density and the applied filtering threshold used to generate the final, high-confidence datasets for model training and evaluation. *Note that most proteins are dissimilar and thus do not reach a method specific threshold, i*.*e. they can not be included into the set visualized herein*. **(A) Sequence-based metrics**. Distributions derived from *the* MMSeqs2 search. The panels show, from left to right: the minimum alignment coverage between sequence pairs, the fractional sequence identity (fident), and the calculated HFSP scores. **(B) Structural confidence metric**. Distribution of the average per-residue confidence scores (pLDDT) for all protein structures from the AlphaFold DB (v4). **(C) Structural similarity metrics**. Distributions derived from Foldseek search. The panels show the minimum structural alignment coverage and the resulting alignment TM-score (alntmscore). In each panel, the red dashed line indicates the specific filtering threshold applied. The red shaded area highlights the portion of the data that was excluded based on this threshold. The associated text annotation quantifies the absolute number and corresponding percentage of protein pairs (or individual structures in B) that were removed by each filter.

**Figure S2:**
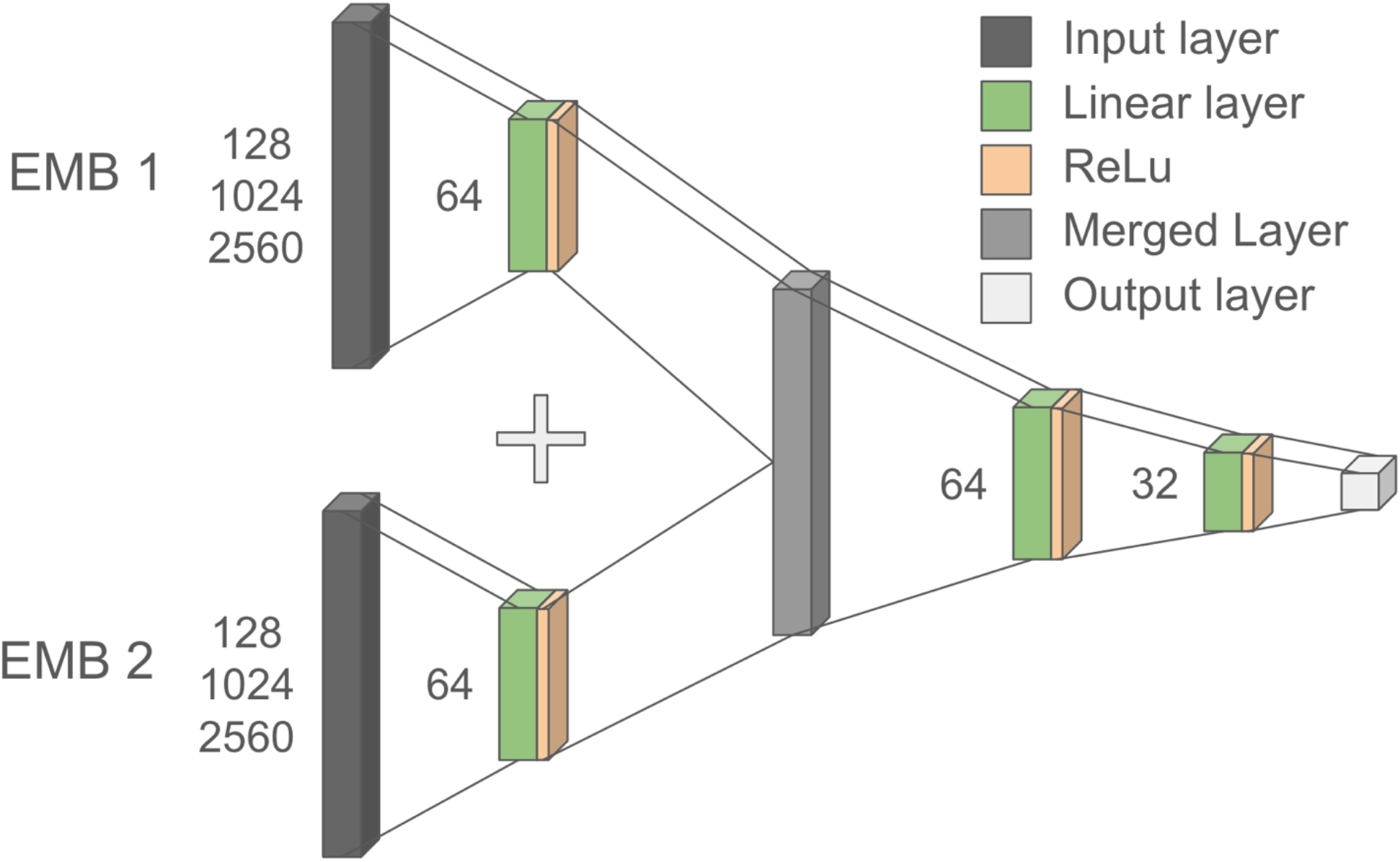
Feed-forward network architecture for supervised learning on protein pair relationships. The network processes two protein embeddings (EMB1 and EMB2) of dimension n (pLM-dependent) through a symmetric architecture. Each embedding passes through a shared weight layer (n → 64; shown separately for visualization), followed by concatenation and successive fully connected layers (128 → 64 → 32 → 1). The output represents the predicted biological relationship between the protein pair (sequence identity, structural similarity, or functional similarity).

**Figure S3:**
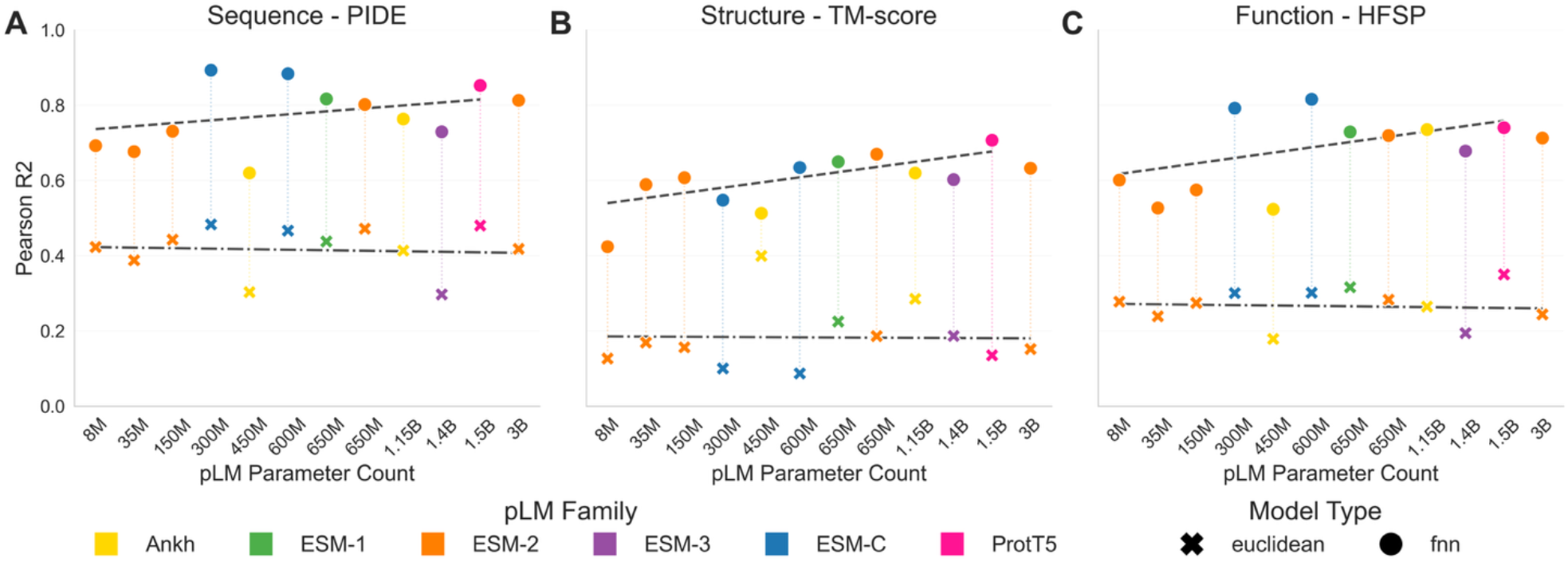
Performance with PCA-compressed embeddings mirrors the trends in Figure 1. Each panel reports the Pearson R^2^ on SwissProt-pre2024 test pairs for (A) sequence identity (PIDE), (B) structural similarity (alignment TM-score), and (C) functional similarity (HFSP). The x-axis lists twelve non-specialist pLMs ordered by parameter count within each family; colors encode families (Ankh = yellow, ESM-1 = green, ESM-2 = orange, ESM-3 = purple, ESM-C = blue, ProtT5 = pink). For every model, a × marks performance from the Euclidean distance between PCA-projected embeddings (inherent information after compression), while a • shows an FNN trained on the same PCA features (extractable information after compression). Points within a family are linked by faint dotted lines. Error bars are not visible since standard errors are all below 0.001. As in Figure 1 trendlines across pLM parameter sizes (dashed line for FNN, dash-dot line for Euclidean) show that inherent signal (×) plateaus with model size, with slopes of -0.0051, -0.0017, and -0.004 for panels A, B, and C respectively, while extractable information (•) increases, with slopes 0.053, 0.092, and 0.095 (all values in change per billion parameters). Note: ESM2-3B was excluded from the FNN trendline fit.

**Figure S4:**
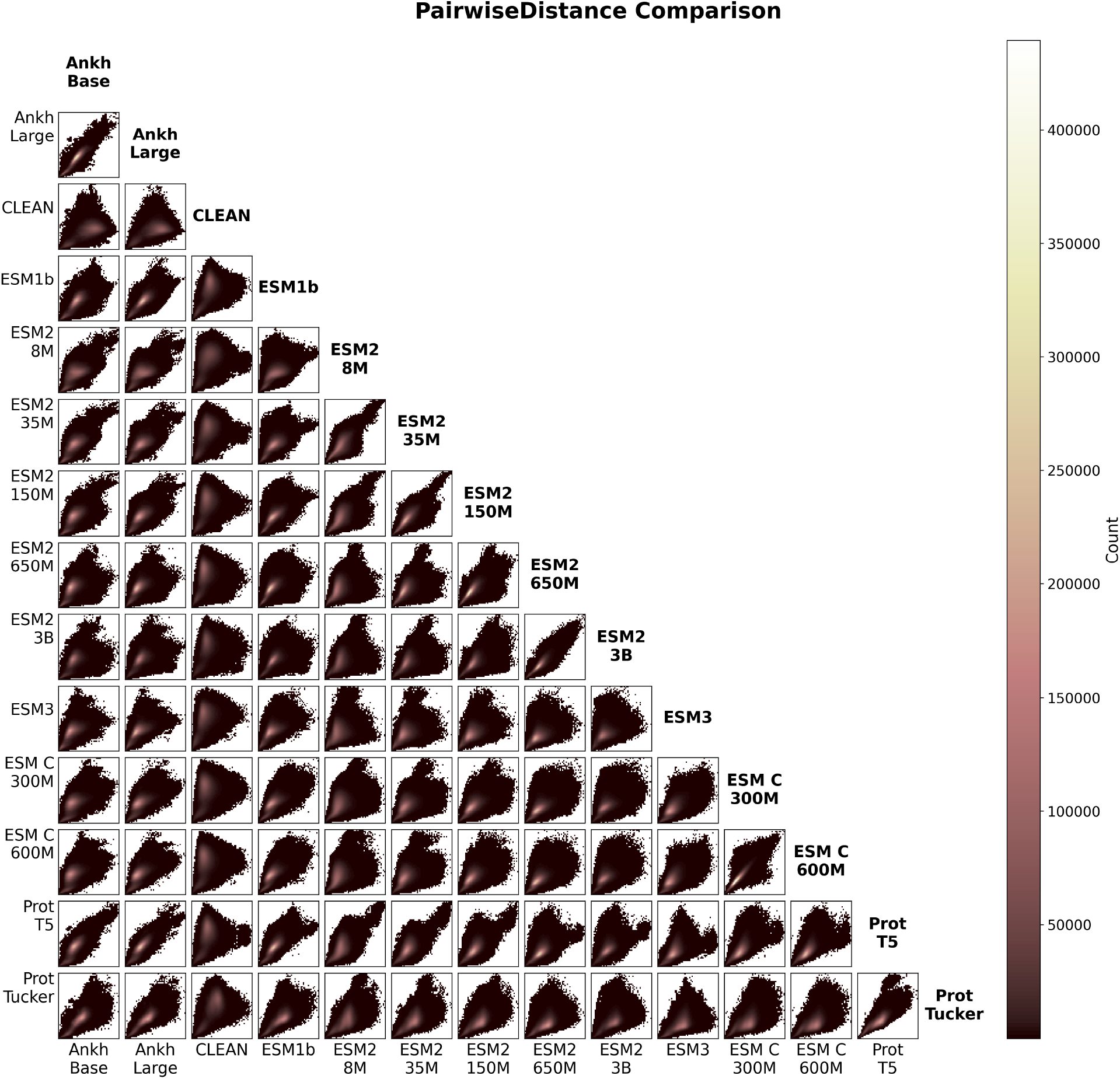
pLM distance distributions reveal family-specific embedding geometries through density fingerprints. Hexagonal grid plots comparing pairwise Euclidean distance distributions across all pLM pairs in the SwissProt-pre2024 dataset. For every model pair, we plot the joint distribution of distances using a 50 × 50 hexagonal grid, where color intensity represents the number of protein pairs falling in each matched distance bin. Each grid cell visualizes how distance values from one model align with those from another across all protein pairs. The diagonal displays model names for orientation. Tight, concentrated distributions along the diagonal indicate similar distance geometries between models, while dispersed patterns reveal divergent embedding spaces. Models within the same family (e.g., Ankh-base/large, ESM-C variants) show concentrated joint distributions, reflecting shared architectural principles. In contrast, CLEAN exhibits dispersed patterns when compared to foundation models, consistent with its unique contrastive training objective.

## Notes

### Competing Interest Statement

The authors have declared no competing interest.

